# Benchmarking BarraCUDA on Epigenetic DNA and nVidia Pascal GPUs

**DOI:** 10.1101/095075

**Authors:** W. B. Langdon

## Abstract

Typically BarraCUDA uses CUDA graphics cards to map DNA reads to the human genome. Previously its software source code was genetically miproved for short paired end next generation sequences. On longer, 150 base paired end noisy Cambridge Epigsnetix’s data, a Pascal GTX 1080 proc esses about 10000 strings per second, comparable with twin nVidia Tesla K40.

## 1 Introduction

The green lines in Figure 1 show graphics processing (GPU) speeds have continued to outstrip conventional computing (CPU blue). This, has prompted interest in using highly parallel graphics cards for general computing (GPGPU computing [7]) and in particular for biological computing [3].

**Figure 1:**
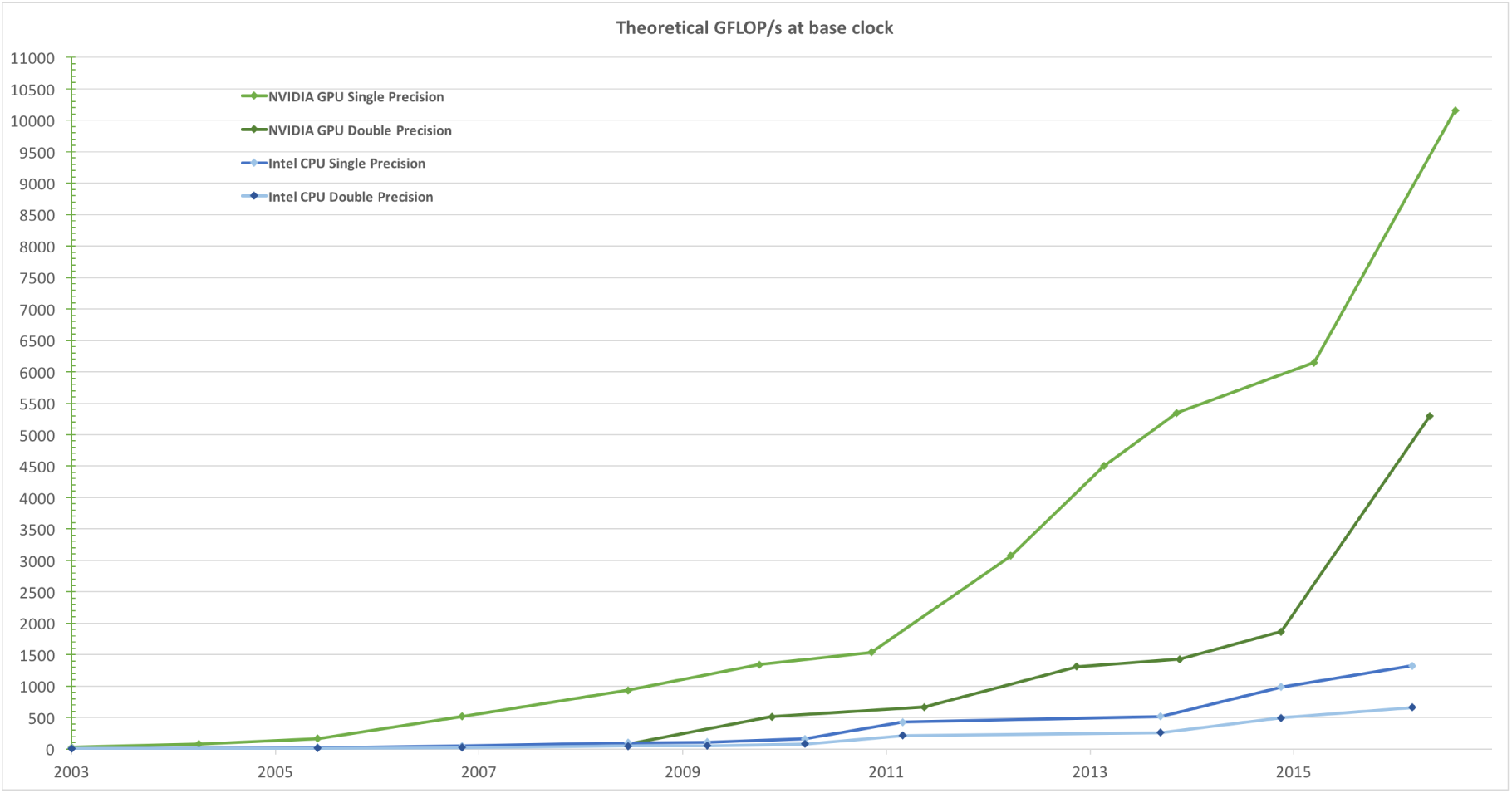
Historical trends in CPU and GPU speeds (nVidia’s CUDA 8 C Programming Guide, Figure 1.)

We have already measured the speed of the GI [2] version of [BarraCUDA](http://seqbarracuda.sourceforge.net/) [1], a popular Bioinformatics tool, on a GCAT next generation DNA sequence benchmark [4] and an epigenetics benchmark provided by Cambridge Epigenetix [6]. Here we re-run the CEGX tests on more recent and more powerful nVidia graphics hardware. The interested reader is referred to [4, 6] for more background.

The GI version of [BarraCUDA](http://seqbarracuda.sourceforge.net/) [5] has been downloaded from SourceForge more than 4000 times since it was released. It was optimised for aligning short next generation (NGS) DNA reads to the reference human genome. Typically epigenetics data uses a reference index file which is approximately twice as big as that used with NGS DNA sequences. Fortunately, with the advent of nVidia Tesla K40 and Titan GPUs, it is now possible to place epigenetics reference datasets onto GPUs. (Appendix C describes the minor change needed to BarraCUDA.)

The human genome, in particular, contains many repeated sequences. This means in many cases a single relatively short DNA sequence may match the reference genome in many places. However the NGS technology is not well suited to obtaining much longer sequences. Therefore it is common to minimise the problem of repeated matches by sequencing both ends of a much longer DNA strand. This leads to “paired end” NGS data, where the sequencing tools (such as BarraCUDA) align each end separately. Due to the repeated sequences, each end may match multiple times, but knowing approximately how far about the two ends are makes it possible to discard many repeated matches as being too far apart to be feasible. BarraCUDA uses two separate “aln” processes, whose output is fed into a “sampe” [SAM paired-end] process which winnows the multiple aln matches and produces the best pair of matches in SAM format.

Since only one GPU of each type was available, we did not attempt to repeat the dual alignment with unix pipes (described in [6]). Instead each step was done sequentially. That is the BarraCUDA aln was run on the R1 DNA data and the results stored to a local temporary binary .sai file. This was immediately followed by BarraCUDA aln on the R2 DNA data. Finally BarraCUDA sampe processed both .sai files to generate the best combined alignment in plain text SAM format.

## 2 CUDA 8.0 Epigentics Results

BarraCUDA was run on a 32 core 2.40GHz UCL cluster node with 126 GB of RAM with CUDA version 8.0 installed on CentOS 6.5 and using version 4.4.7 of the GNU gcc compiler. It housed a TITAN X (Pascal), a GeForce GTX 1080 and a GeForce GTX TITAN X (see Table 1).

**Table 1:**
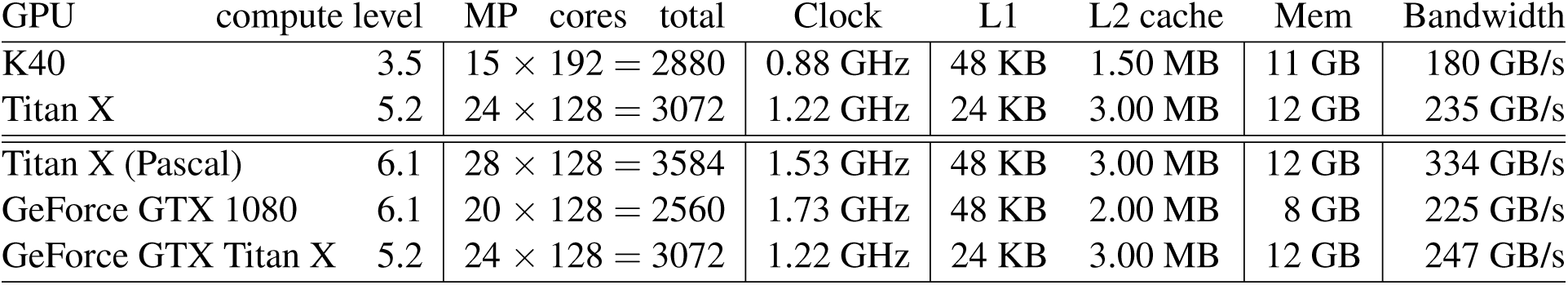
GPU Hardware. Second column is CUDA compute capability level (as can be used with nvcc’s `-arch`). Each GPU chip contains 15, 20, 24 or 28 identical independent multiprocessors (MP, column 3). Each MP contains 192 or 128 stream processors (total given in column 5), whose clock speed is given in column 6. The next two columns give sizes of the L1 and L2 caches used. The on chip cache sizes are followed by (off chip) on board memory and, in the last column, the maximum measured data rate between the GPU and its on board memory. First rows are for comparison only [6].

The epigenetics data are considerably noisier than the GCAT NGS data. The usual default for BarraCUDA is to allow only two mismatches. With noisier data, it must be instructed (via the `–n` switch) to allow more mismatches. It takes more searching to find strings which diverge from the reference genome in more places so slowing down `aln`. Also typically noise increases the number of potential approximate matches so causing more work for the `sampe` process. Although BarraCUDA `sampe` can use multiple CPU threads, eventually additional threads give little additional benefit.

With paired epigenetics reads, the R1 and R2 strands of DNA typically have different amounts of noise. This means one of the `aln` processes has much more work to do than the other. (See the separation of the R1 and R2 `aln` plots in Figure 2.) Although the new server has 32 cores, they each run more slowly than the 12 core server we previously used [6] (plotted in black in Figure 2). The sampe process gains little from using the additional threads and the speed ratio on the two servers, 1.35 on average, is close to the ratio of the two servers’ CPU clock frequencies.

**Figure 2:**
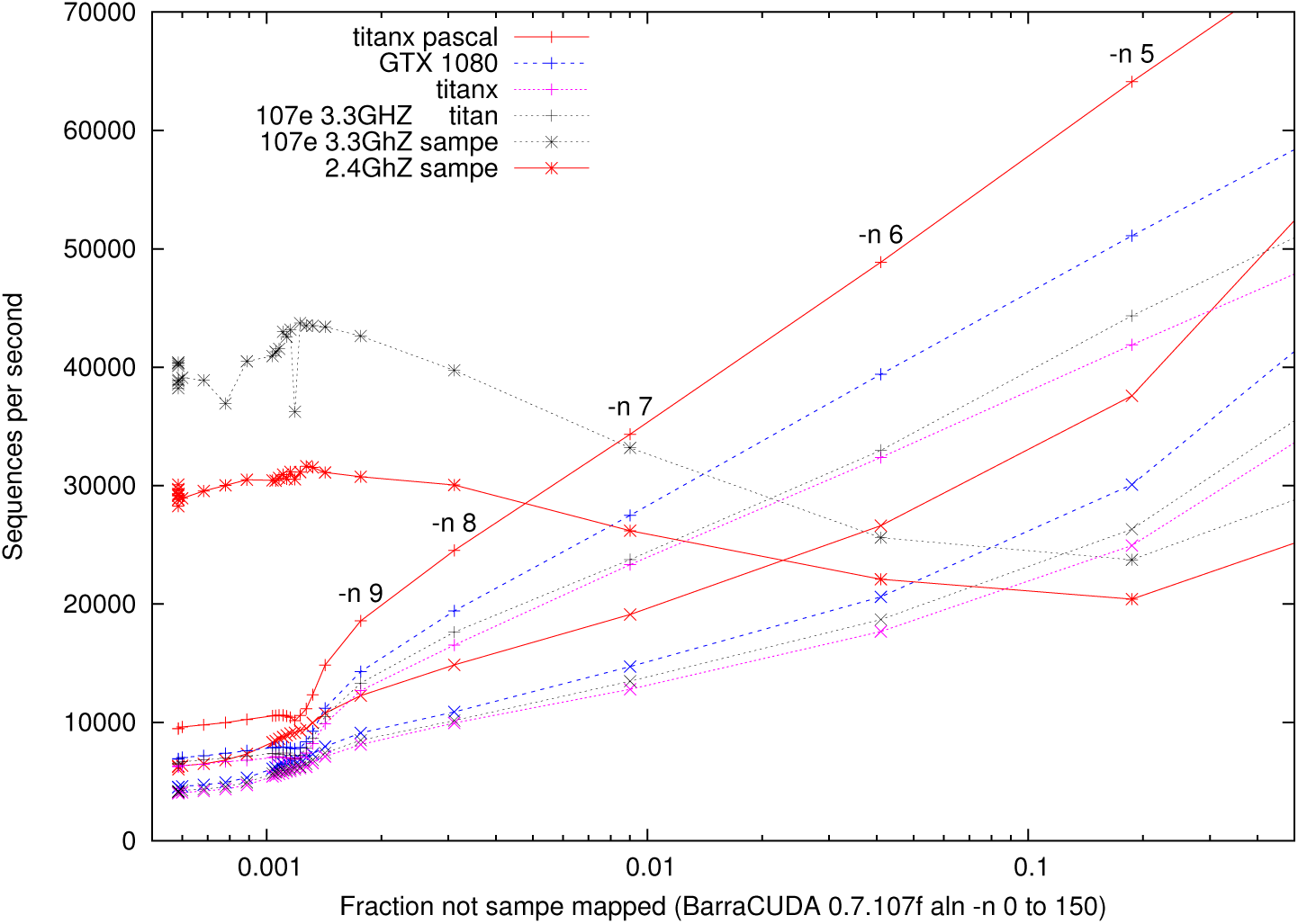
Speed of BarraCUDA on 10 million simulated 150 base pairs epigenetics paired end reads. Solid lines are the speed of BarraCUDA alignment on an nVidia 12 GByte Titan X (pascal, compute level 6.1). (Asymmetry between the R1 + and R2 × ends is due the expected difference in noise on the R1 and R2 data.) GTX 1080 and Titan X (See Table 1) plotted with dashed blue and dotted purple lines.) Speed of sampe process * is dominated by the CPU clock speed. (Sampe does not use the GPU.) For comparison, old data plotted in black.

With the default number of mismatches (`–n 2`) `aln` is blisteringly fast but almost no matches are found. `-n` must be increased to 7 to get reasonable results, whereupon the Titan X (pascal) processes 34000 sequences per second (R1), 19 000 (R2) and the 32 core CPU sampe processes outputs 28 000 answers per second. Assuming copious data and reasonable overlapping of both aln and sampe processes, a single GPU is limited by the slowest step (i.e. aln R2). Nonetheless, depending upon noise, at -n 7, a single Titan X (pascal) should process about 12 000 150BP paired end sequences per second. Similarly, for a single GTX 1080 the figures are: 27 000 (R1), 15 000 (R2), 26 000 (sampe), 10 000 (combined). For the Titan X: 23 000 (R1), 13, 000 (R2), 26000 (sampe), 8 000 (combined) SAM answers per second. (Making similar assumptions for the earlier experiments with CUDA 6 Barracuda 0.7.107e that Titan card (see Table 1) on a 12 core CPU server gave 24000 (R1) 13 000 (R2), 33 000 (sampe). Giving an estimated total of 8 000 SAM answers per second. Whereas for the pair of K40s, on a different server but with real overlap, gave 15 000 (R1), 8 600 (R2), 7 900 (sampe) and a total of 7 700 sequences per second.)

To summarise. Using `-n` 7 (to get a match rate above 99%), with copious paired end CEGX data, with a single GPU on a many core CPU, BarraCUDA sampe can be run in parallel with the aln processes so that it appears to impose no overhead. Instead BarraCUDA is limited by the two aln steps. Thus we should get total throughput of about:

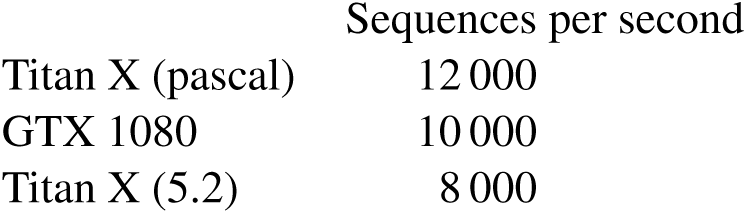

|  | Sequences per second |
| --- | --- |
| Titan X (pascal) | 12 000 |
| GTX 1080 | 10 000 |
| Titan X (5.2) | 8 000 |

## Acknowledgements

I would like to thank Tristan Clark, Justyna Petke and njuffa.

CEGX DNA methylation sequences supplied by Albert Vilella Cambridge Epigenetix

## A Epigenetics Fasta Sequences

Cambridge Epigenetix generated ten million pairs of synthetic epigenetics data, each end comprising 150 base pairs. Data available via: https://s3-eu-west-1.amazonaws.com/cegx-test-001/tmwg-example-files/SIM03_S1_L001_R1_001.fastq.gz and https://s3-eu-west-1.amazonaws.com/cegx-test-001/tmwg-example-files/SIM03_S1_L001_R2_001.fastq.gz.

## B Epigenetic (HS) Human Reference Genome

BarraCUDA was used to build its index file from the Epigenetics fasta sequences generated from the reference human genome (38DH). It (hs38DH) is 6 Gbytes. I.e. about twice the size of usual BarraCUDA index. It may be down loaded from https://s3-eu-west-1.amazonaws.com/cegx-test-001/tmwg-example-files/hs38DH_bwameth.tar.gz.

## C Code Changes to BarraCUDA

There was a problem in version 0.7.107f on the 1080 which made the main buffer too large. It was halved in size by hand.

## References

[1] P. Klus, S. Lam, D. Lyberg, M. S. Cheung, G. Pullan, I. McFarlane, G. S. H. Yeo, and Brian Y. H. Lam. BarraCUDA - a fast short read sequence aligner using graphics processing units. BMC Research Notes, 5(27), 2012.

[2] W. B. Langdon. Genetically improved software. In A. H. Gandomi et al., editors, Handbook of Genetic Programming Applications, chapter 8, pages 181–220. Springer, 2015.

[3] W. B. Langdon. Computational biology in the 21st century is parallel. Communications of the ACM, 59(11):9, November 2016. Letter to the editor.

[4] W. B. Langdon and Brian Y. H. Lam. Genetically improved BarraCUDA. Research Note RN/15/03, Department of Computer Science, University College London, Gower Street, London WC1E 6BT, UK, 28 May 2015.

[5] W. B. Langdon, Brian Y. H. Lam, J. Petke, and M. Harman. Improving CUDA DNA analysis software with genetic programming. In S. Silva et al., editors, GECCO, pages 1063–1070, Madrid, 11-15 July 2015. ACM.

[6] W. B. Langdon, A. Vilella, Brian Y. H. Lam, J. Petke, and M. Harman. Benchmarking genetically improved BarraCUDA on epigenetic methylation NGS datasets and nVidia GPUs. In J. Petke et al., editors, Genetic Improvement 2016 Workshop, pages 1131–1132, Denver, July 20-24 2016. ACM.

[7] J. D. Owens, M. Houston, D. Luebke, S. Green, J. E. Stone, and J. C. Phillips. GPU computing. Proceedings of the IEEE, 96(5):879–899, May 2008. Invited paper.

